# Selective amplification of ipRGC signals accounts for interictal photophobia in migraine

**DOI:** 10.1101/2020.04.17.047290

**Authors:** Harrison McAdams, Eric A Kaiser, Aleksandra Igdalova, Edda B Haggerty, Brett Cucchiara, David H Brainard, Geoffrey K Aguirre

## Abstract

Second only to headache, photophobia is the most debilitating symptom reported by people with migraine. While the melanopsin-containing, intrinsically photosensitive retinal ganglion cells (ipRGCs) are thought to play a role, how cone and melanopsin signals are integrated in this pathway to produce visual discomfort is poorly understood.

We studied 60 people: 20 without headache and 20 each with interictal photophobia from migraine with or without aura. Participants viewed pulses of spectral change that selectively targeted melanopsin, the cones, or both, and rated the degree of visual discomfort produced by these stimuli while we also recorded pupil responses.

We examined the data within a model that describes how cone and melanopsin signals are weighted and combined at the level of the retina, and how this combined signal is transformed into a rating of discomfort or pupil response. Our results indicate that people with migraine do not differ from headache-free controls in the manner in which melanopsin and cone signals are combined. Instead, people with migraine demonstrate an amplification of integrated ipRGC signals for discomfort. This effect of migraine is selective for ratings of visual discomfort, in that an amplification of pupil responses was not seen in the migraine group, nor were group differences found in surveys of other behaviors putatively linked to ipRGC function (chronotype, seasonal sensitivity, presence of a photic sneeze reflex).

By revealing a dissociation in the amplification of discomfort versus pupil response, our findings suggest a post-retinal alteration in processing of ipRGC signals for photophobia in migraine.

**Significance:** The melanopsin-containing, intrinsically photosensitive retinal ganglion cells (ipRGCs) may contribute to photophobia in migraine. We measured visual discomfort and pupil responses to cone and melanopsin stimulation—the photoreceptor inputs to the ipRGCs—in people with and without migraine. We find that people with migraine do not differ from those without headaches in how cone and melanopsin signals are weighted and combined to produce visual discomfort. Instead, migraine is associated with an amplification of ipRGC signals for discomfort. This effect of migraine upon ipRGC signals is limited to photophobia, as we did not find an enhancement of pupil responses or a change in other behaviors linked to ipRGC function. Our findings suggest a post-retinal amplification of ipRGC signals for photophobia in migraine.

## Introduction

People find bright light uncomfortable and sometimes even painful. This experience of light-induced discomfort is exacerbated in numerous clinical conditions and can be debilitating^1^. We refer here to discomfort from light as photophobia, which is typically manifest as a somatic sensation localized to the eyes or head^2^. A common cause of photophobia is migraine^3^. Photophobia is reported by 80-90% of individuals during a migraine attack^4–6^, and 50% of individuals report it as their most burdensome symptom^7^. Even between headaches, people with migraine have a lowered threshold for pain from light as compared to headache-free controls^8–11^.

The signals that ultimately result in photophobia presumably begin with photoreceptors in the eye. Under daylight conditions, the cone photoreceptors capture photons and relay signals via retinal ganglion cells to thalamic and brainstem targets (Figure 1a). A subset of retinal ganglion cells express the photopigment melanopsin^12^. These “intrinsically photosensitive” retinal ganglion cells (ipRGCs) are capable of responding to light without synaptic input^13^. There is evidence from rodent studies that ipRGCs project to the somatosensory thalamus, where they innervate neurons that are also sensitive to dural stimulation carried by trigeminal afferents^14^ (Figure 1a). This finding offers a neural mechanism by which light stimulation creates somatic discomfort. The ipRGCs contribute to other “reflexive” functions of vision as well, including photo-entrainment of the circadian rhythm^15,16^ and control of pupil size^17–19^.

**Figure 1.**
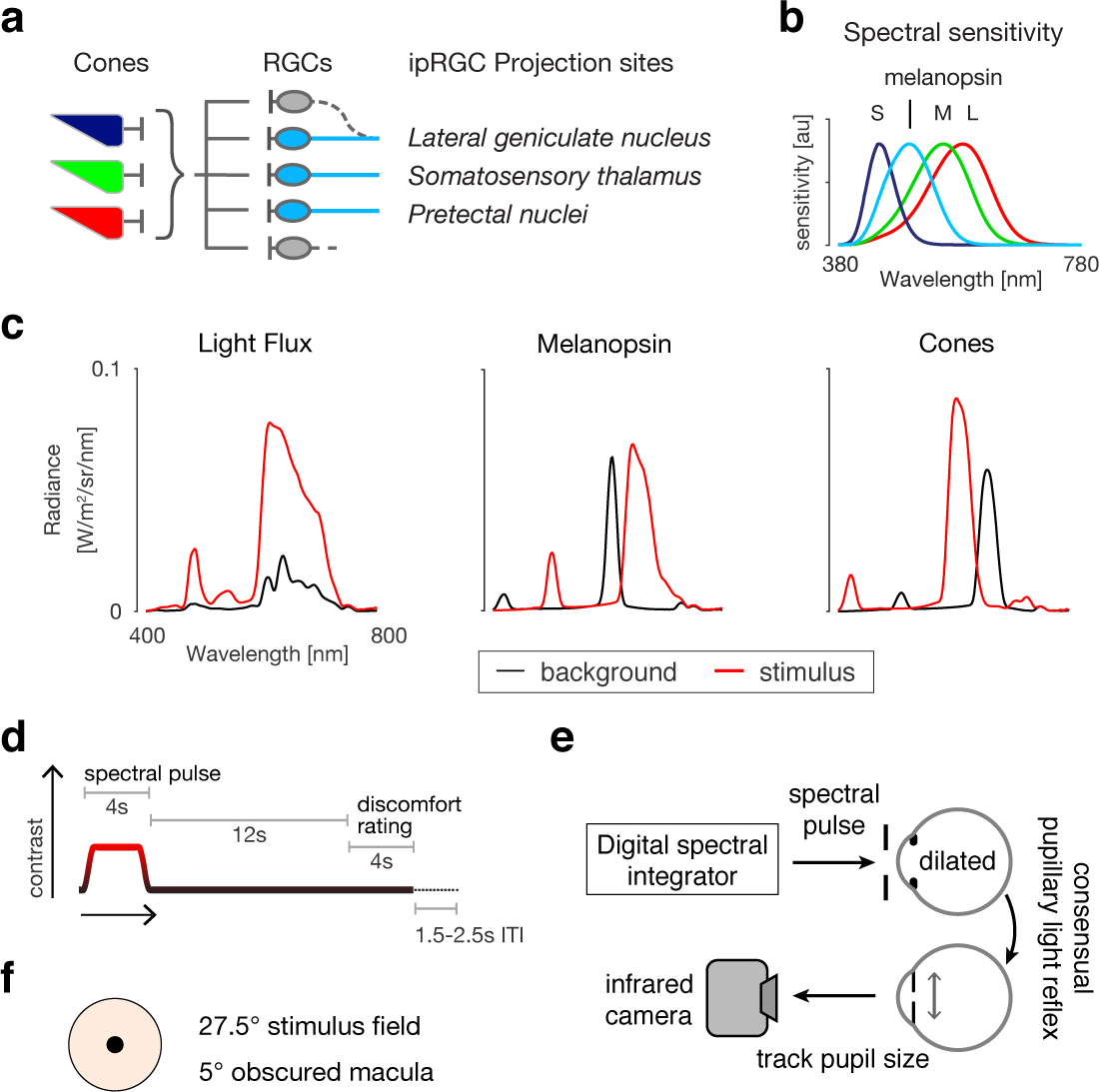
Experiment Overview. **a**. There are several classes of melanopsin-containing ipRGCs which vary in their central projections, function, and extent to which they receive input from cones. Among other sites, the ipRGCs project to the somatosensory thalamus and the lateral geniculate nucleus, where their signals may contribute to light sensitivity. Other ipRGCs project to the pretectal nuclei to control the size of the pupil. **b**. The spectral sensitivity functions of the relevant photoreceptors under daylight conditions. **c**. Shown are pairs of spectra (background: black; stimulus: red) that differ in excitation for the targeted photoreceptors. From left to right, the stimuli produce: equal contrast on the cones and melanopsin (termed light flux); contrast only on melanopsin; and equal contrast across all three classes of cones but no contrast on melanopsin. **d**. Each trial featured a four-second period during which the stimulus transitioned from the background to the stimulation spectrum and back. Twelve seconds after stimulus offset, the subject provided a discomfort rating. There was an inter-trial interval that varied between 1.5 and 2.5 s. **e**. The light from a digital spectral integrator was presented to the pharmacologically dilated right eye of the subject through an artificial pupil. The consensual pupillary light response of left eye was recorded with an infrared camera. **f**. The stimulus spectra were presented in an eyepiece with a 27.5 degree diameter field, with the central 5 degrees obscured to minimize macular stimulation.

The ipRGCs may play a role in human photophobia. People who have migraine and are also blind from inherited rod-cone degeneration experience photophobia during a headache^14^, implicating spared ipRGCs as the source of this sensation. In people without visual impairment, Stringham and colleagues found that shorter wavelengths of light (closer to the peak spectral sensitivity of the melanopsin photopigment) tend to produce greater discomfort in healthy observers^20^. Studies that use narrow-band light stimuli, however, are limited in their ability to probe the specific contribution of melanopsin to photophobia in the intact visual system. This is due to the considerable overlap of the cone and melanopsin spectral sensitivity functions (Figure 1b). Moreover, some classes of ipRGCs also receive input from the cones^13,21–23^. As a consequence, photophobia may result from both melanopsin and cone signals after their integration within ipRGCs. It is unknown how these photoreceptor classes are weighted and combined to produce photophobia, and how this process might be altered in migraine.

In previous work we have shown that carefully tailored modulations of the spectral content of light may be used to selectively target melanopsin or the cones.^24,25^ Here, we examine the contribution of cone and melanopsin signals to visual discomfort, and to pupil responses, in people who have migraine with interictal photophobia. Participants reported the discomfort they experienced from viewing pulses of light that selectively targeted melanopsin, the cones, or their combination (Figure 1c, d). Pupillometry in response to these pulses was also obtained (Figure 1e). We recruited 20 participants in each of three groups: migraine with visual aura, migraine without aura, and headache-free controls. All of the participants with migraine endorsed interictal sensitivity to light. Our findings demonstrate that both melanopsin and cone stimulation in isolation produce visual discomfort. By examining the effect of separate and simultaneous stimulation of melanopsin and the cones, we quantified how these photoreceptor signals are weighted and combined to produce visual discomfort and pupil responses. We find that the enhanced interictal light sensitivity observed in migraine is well described as an amplification of photoreceptor signals after their combination. We further demonstrate that pupil responses are governed by different combination parameters, and do not demonstrate amplification in migraine. These results indicate that interictal photophobia in migraine is a selective amplification of a subset of ipRGC outputs, most plausibly at a post-retinal locus.

## Results

### Participant demographic and clinical characteristics

We studied 20 people from each of three groups: migraine with aura (MwA), without (MwoA) aura, and headache free controls (HAf). The three groups (Table 1) were well-matched in age (F[2,57] = 0.2, p = 0.820), but differed in gender distribution (F[2,57] = 3.3, p = 0.0439), with fewer women in the control group. The greater proportion of women in the migraine groups is consistent with migraine epidemiology^26^. Headache frequency was similar in the two migraine groups with 12 (± 10) and 13 (± 9) days with headache reported within a 90 day period by MwA and MwoA subjects, respectively (∼4 headache days per month), consistent with a classification of episodic (as opposed to chronic) migraine^27^. Acetaminophen and NSAID use for any indication were similar across all three groups. Triptan use was reported by 5 MwA and 1 MwoA participants. Similarly, combined aspirin/acetaminophen/caffeine use was reported by 5 MwA and 1 MwoA participants. Preventive medication use (e.g., tricyclics, beta-blockers, etc.) was reported by 1 MwA and 3 MwoA participants. We quantified headache disease burden using the MIDAS^28^ and HIT-6^29^ surveys. The migraine groups unsurprisingly had higher scores on both instruments relative to headache-free controls (MIDAS: F[2,57] = 13.65, p = 1.43e-5; HIT-6: F[2,57] = 48.82, p = 4.43e-13). The two migraine groups did not differ in disease impact (MIDAS: t = 1.00, p = 0.76; HIT-6: t = 0.40, p = 0.96). The distribution of these values suggests moderate disability from migraine in both groups.

**Table 1.** Subject demographic and clinical characteristics. Participants were asked to report the number of headaches they had experienced over the prior three months. The Migraine Disability Assessment Test (MIDAS)^28^ and the Headache Impact Test (HIT-6)^29^ measure headache disability. Medication use is summarized within four categories. Where appropriate, the mean value (and standard deviation) across subjects is reported.

| Group | # women | Age in years | Headache Days per 3 Months | Disability |  | Medication use |  |  |  |
| --- | --- | --- | --- | --- | --- | --- | --- | --- | --- |
|  |  |  |  | MIDAS | HIT-6 | NSAID | Excedrin | Triptan | Preventive |
| Controls | 13/20 | 31 (5) | 1.3 (1.4) | 0.5 (0.8) | 40.7 (4.1) | 15 | 0 | 0 | 0 |
| MwA | 19/20 | 31 (4) | 13.1 (8.9) | 18.6 (15.3) | 60.6 (8.0) | 16 | 6 | 5 | 1 |
| MwoA | 17/20 | 30 (4) | 11.7 (9.7) | 16.0 (13.7) | 60.0 (8.8) | 18 | 1 | 1 | 3 |

### Participants with migraine have interictal photophobia, but do not differ from controls in surveys of circadian and seasonal behavior

The Visual Discomfort Scale (VDS) measures symptoms of discomfort from reading, patterns, and light on a 0-69 scale^30^. We required our control participants to have a low score on this instrument (≤ 7) but did not impose a requirement for migraineurs. Symptoms of visual discomfort were correspondingly greater in the migraine population as compared to the controls (Table 2, F[2,57] = 15.23, p = 5.02e-6). Participants also completed the Photosensitivity Assessment Questionnaire (PAQ) which measures light-avoiding (“photophobia”) and light-seeking (“photophilia”) behavior on a 0-8 scale^31^. Migraine participants again reported greater light avoidance as compared to controls (Table 2, F[2,57] = 10.95, p = 9.44e-5), although there was no difference in reported light-seeking behavior (Table 2, F[2,57] = 0.75, p = 0.448).

**Table 2.**
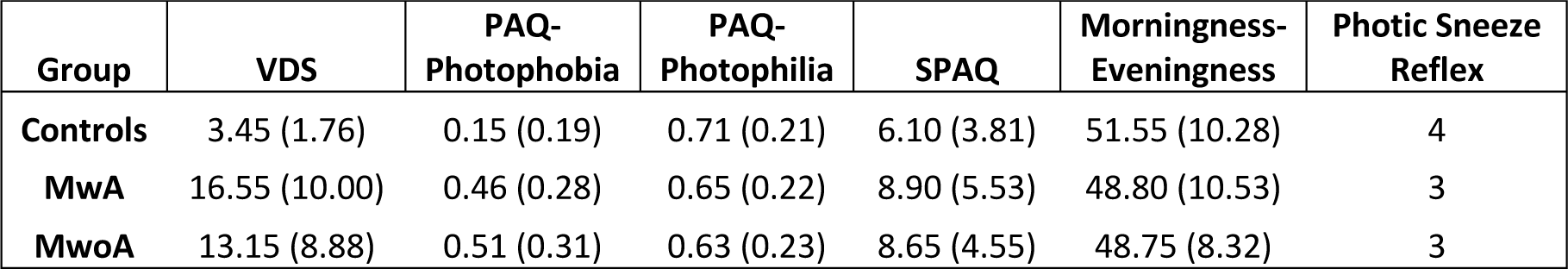
Surveys of behaviors that may be related to ipRGC function. The Visual Discomfort Scale (VDS) measures reported light sensitivity across several domains of visual function^30^. The Photosensitivity Assessment Questionnaire (PAQ) measures reported “photophobia” and “photophilia” behaviors ^31^. The Seasonal Pattern Assessment Questionnaire (SPAQ)^33^ measures the reported degree to which mood and behavior varies over course of a year, and the Morningness-Eveningness Questionnaire provides a “chronotype” score^32^. Values are the mean (and standard deviation) across subjects within each group. Finally, we asked subjects if they “tend to sneeze when [they] step out of a dark room into bright sunlight” and report here the number of subjects in each group who responded “yes”.

As we are interested in how migraine and photophobia may relate to ipRGC function, we examined if our participant groups differed in other functions thought to be mediated by ipRGCs. In the rodent, multiple classes of ipRGCs have been identified that differ in their subcortical projections and in their functional properties. Projections of the ipRGCs to the suprachiasmatic nucleus are thought to control circadian photoentrainment^15^. As variation in this function is speculated to relate to sleep alterations and seasonal affective disorder, we gathered information about the sleep habits and seasonal preferences of our participants (Table 2). The Morningness-Eveningness Questionnaire^32^ characterizes chronotype on a scale of 16-86, with the extremes corresponding to evening and morning preference, respectively. The median scores for the three groups all were in the mid-range (∼50), and were not significantly different (F[2,57] = 0.54, p = 0.586). The Seasonal Pattern Assessment Questionnaire^33^ provides a Global Seasonality Score, which assesses on a 0-24 scale the degree to which mood and physiology varies across seasons; a score of 16 or higher is typical in patients with seasonal affective disorder. The central tendency of our participants (a score of ∼7) indicates a mild degree of seasonal sensitivity, and this did not differ between the groups (F[2,57] = 2.19, p = 0.121). Finally, the photic sneeze reflex has been hypothesized to be related to ipRGC function^34^. We asked our participants if they experience this phenomenon and did not find any difference between groups in the proportion of participants (15-20%) who have this experience (F[2,57] = 0.04, p = 0.962).

Overall, apart from photophobia, our studied populations were well matched in behaviors hypothesized to be related to ipRGC function.

### Melanopsin and cone contrast produce mild discomfort in control participants

Our participants rated the degree of discomfort they experienced while viewing pulses of spectral change that targeted melanopsin, the cones, or combined stimulation of both sets of photoreceptors (termed light flux). The stimuli were designed to increase excitation in the targeted photoreceptor(s) by 100%, 200% or 400%. Participants rated the amount of discomfort produced by each type of light pulse on a 0 (none) to 10 (extreme) scale.

The light flux stimulus combines melanopsin and cone stimulation. In the HAf control participants, light flux pulses evoked mild discomfort, increasing with contrast, reaching a mean discomfort rating of 3.15 out of 10 for 400% contrast (Figure 2, left-top). To determine whether this discomfort was a consequence of melanopsin or cone-based signaling, we examined the discomfort ratings in response to stimuli designed to target these photoreceptor classes in isolation. Discomfort ratings to both melanopsin (Figure 2, left-middle) and cone-directed stimuli (Figure 2, left-bottom) also increased with contrast, but only with mild discomfort at 400% (Figure 2, center row, left column: mean rating of 2.18 for melanopsin; bottom row, left column: 2.80 for cones). This result suggests that both cone and melanopsin signals contribute to light-induced discomfort. For all stimuli, we further observed that logarithmic changes in stimulus contrast produced linear changes in mean rated discomfort, as illustrated by the good agreement between the fit lines and the data (Figure 2).

**Figure 2.**
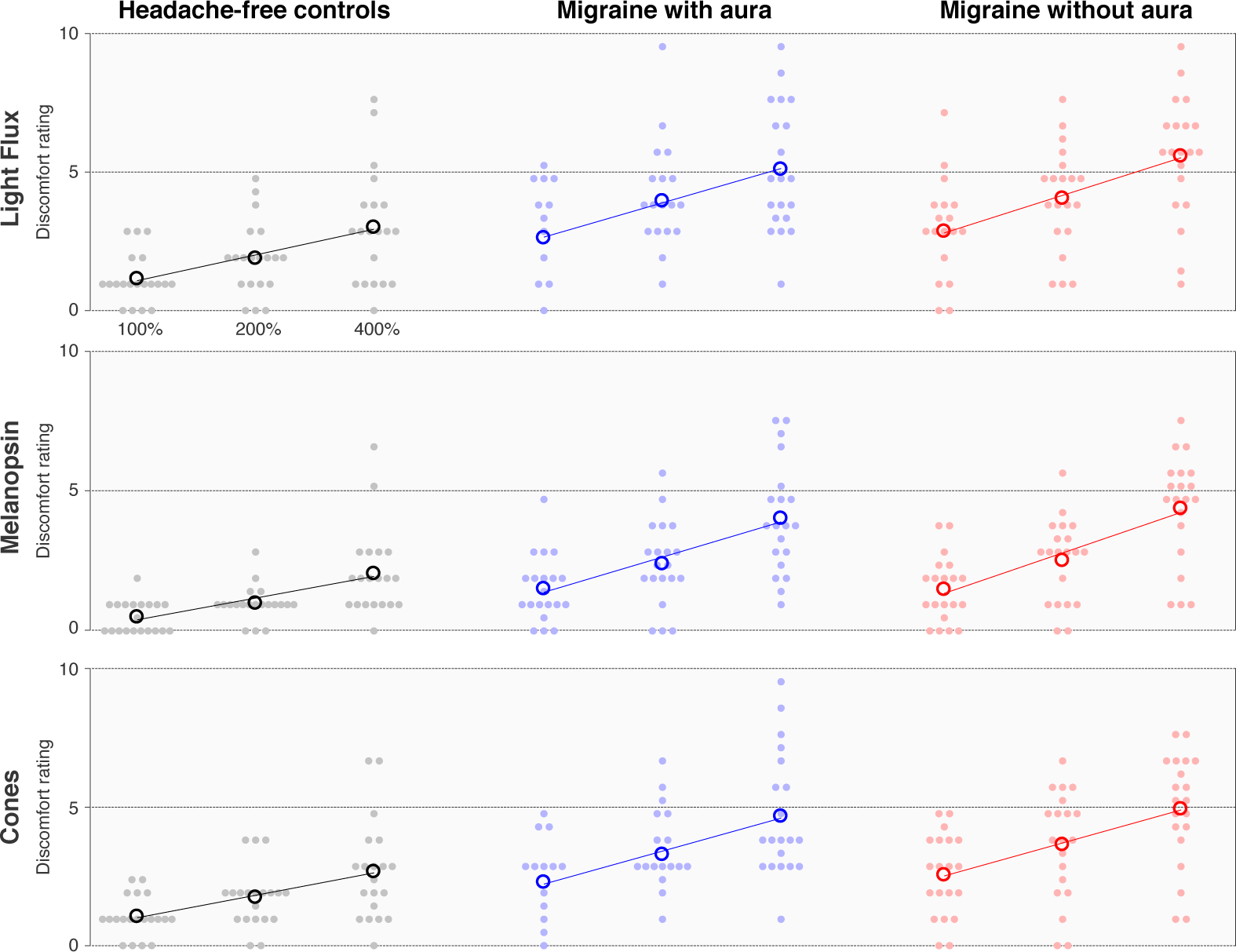
Discomfort ratings by stimulus and group. Each row presents the discomfort ratings elicited by stimuli that targeted a particular combination of photoreceptors, and each column contains the data from each individual group (n = 20 participants per group). The stimuli were presented at three different contrast levels (100, 200, and 400%), and these (log-spaced) values define the x-axis of each subplot. The median (across trial) discomfort rating for a given stimulus and contrast is shown for each participant (filled circle), as is the mean rating across participants (open circle). The best fit line to the mean discomfort rating across participants as a function of log contrast is shown in each subplot.

### Cone and melanopsin signals contribute to interictal photophobia in migraine

We next asked if people with photophobic migraine would experience greater discomfort in response to our stimuli, and if so, whether the enhanced discomfort signal is attributable to the cones, melanopsin, or both. Both migraine groups showed increased discomfort in response to the combined light flux stimuli at all contrast levels (Figure 2, center and right, top: at 400% contrast, mean of 5.35 for MwA and 5.85 for MwoA vs. 3.15 for controls). The mean rating across participants was also increased in both migraine groups in response to melanopsin-directed stimulation (Figure 2, middle row: at 400% contrast, mean of 4.28 for MwA and 4.65 for MwoA vs. 2.18 for controls) and cone-directed stimulation (Figure 2, bottom row: at 400% contrast, mean of 4.90 for MwA and 5.18 for MwoA vs. 2.80 for controls). Both migraine groups also showed a linear relationship between log-spaced contrast and mean discomfort ratings for all stimulus types, which is again illustrated by the fit lines (Figure 2).

There was a higher proportion of women in the migraine groups as compared to the control group. We considered if this unequal distribution of gender could account for the differences in discomfort ratings between the groups. The mean discomfort rating reported by female control participants (across all stimuli) was not higher than the ratings provided by male participants (mean rating men: 1.79, women: 1.74), indicating that differing gender ratios do not account for the increased discomfort in the migraine groups.

### Discomfort ratings are well fit by a two-stage, non-linear, log-linear model

We observe that mean discomfort ratings for all stimuli are well described as a linear function of log-scaled stimulus contrast, consistent with the Weber–Fechner law of perception. It is also apparent that a light flux stimulus, which combines melanopsin and cone contrast, evokes less discomfort than would be predicted given the discomfort produced by each stimulus component alone. These properties of the data may be explained by non-linear combination of melanopsin and cone signals prior to the stage at which photoreceptor signals are interpreted as discomfort.

We examined these impressions within the context of a quantitative, two-stage model governed by four parameters. The first stage is based upon psychophysical measures of combined stimulus dimensions^35,36^, and the second on the Weber-Fechner Law. The model provides a discomfort rating for stimuli with arbitrary combinations of melanopsin and cone contrast.

The first stage of the model (Figure 3a, left) considers the combination of melanopsin and cone signals within ipRGCs. The inputs to this stage are the contrasts on the melanopsin and cone photoreceptors created by a stimulus. A light flux stimulus of (e.g.) 200% contrast has the property of providing 200% contrast input on both of these photoreceptor classes. A scaling factor (α) adjusts the relative potency of melanopsin contrast as compared to cone contrast. The two contrast types are then combined using a Minkowski distance metric with exponent β. This integrated, “ipRGC contrast” is log-transformed and then passed to the second stage of the model (Figure 3a, right), which transforms input into a discomfort rating under the control of a slope and offset parameter (which is the intercept transformed to describe the modeled response to 200% contrast).

**Figure 3.**
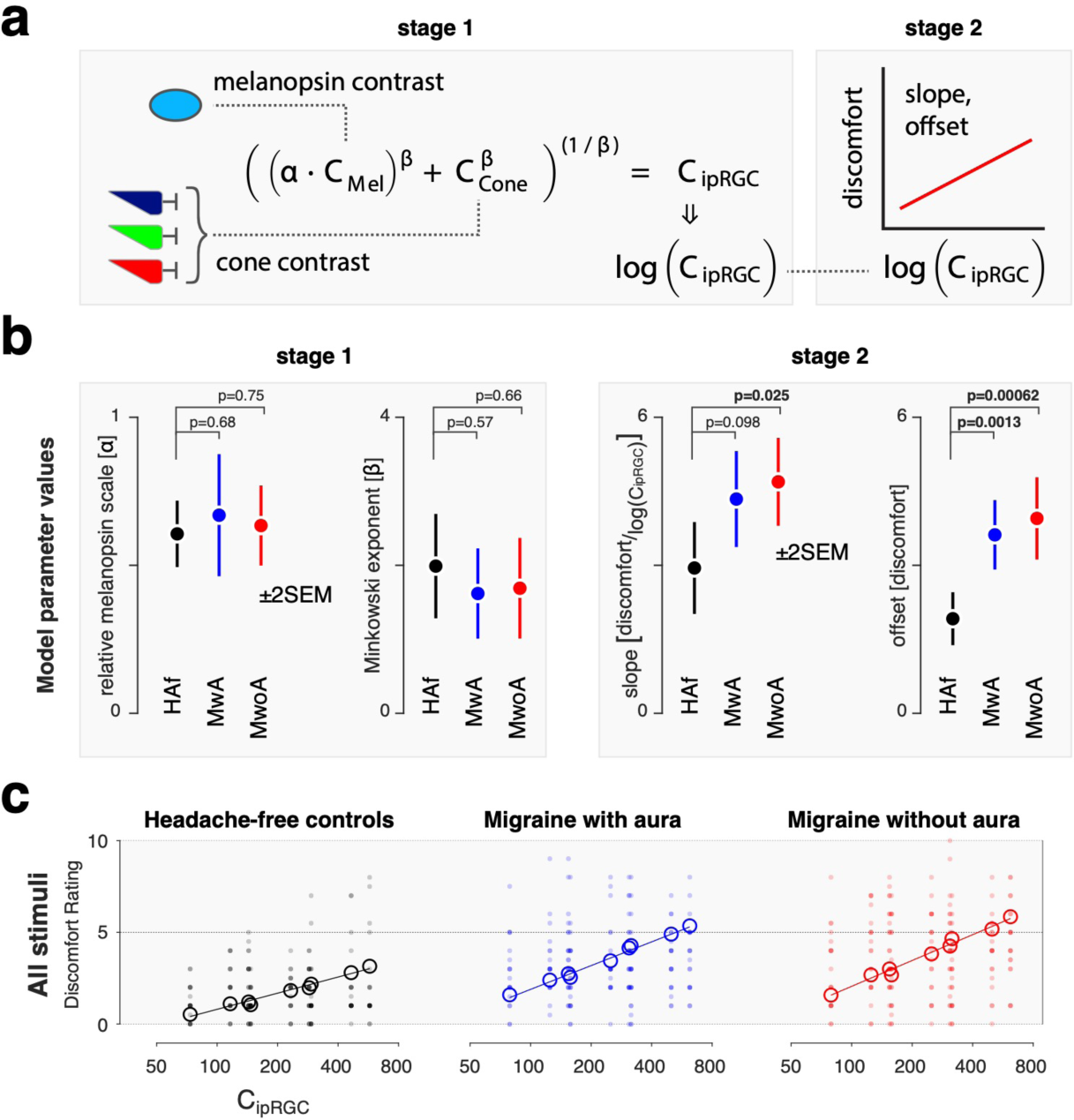
A two-stage model of discomfort ratings. We developed a two-stage model that describes discomfort ratings as a function of melanopsin and cone stimulation. **a**. In the first stage, (left) melanopsin contrast (C_Mel_) is weighted by a scaling factor (α) and then combined with cone contrast (C_Cone_) under the control of the Minkoswki exponent (β). The output of this stage is “ipRGC contrast”, which is log-transformed and passed to the second stage (right). Here, the signal undergoes an affine transform to produce a discomfort rating, under the control of a slope and offset parameter (the latter being expressed as the modelled discomfort rating at 200% ipRGC contrast). **b**. The model was fit to the discomfort data from each group, yielding estimates of the model parameters (±2SEM obtained via bootstrapping). The p-value associated with a two-tailed t-value, taken with respect to the pooled standard errors, is presented for the comparison of each of the migraine groups to the control group for each parameter (n = 20 participants per group). **c**. Stage 1 of the model transforms the stimuli used in the experiment to common units of ipRGC contrast. Each plot presents the discomfort ratings (individual participants in filled circles, group means in open circles) in terms of ipRGC contrast, with the parameters at stage 1 forced to be the same across groups. The fit of the second stage of the model (which can vary across groups) provides the fit line.

We fit this model to the discomfort ratings across trials for all stimuli and participants within a particular group, using bootstrap resampling across participants to characterize the variability of the model parameters. Fitting was performed separately for the data from each group (Figure 3b). The model performed equally well for each group in accounting for the mean (across participant) discomfort ratings across stimuli (model R-squared, ±SEM: HAf controls: 0.95 ± 0.03; MwA: 0.96 ± 0.03; MwoA: 0.97 ± 0.01).

### Migraine groups differ from headache-free controls in the response to integrated melanopsin and cone signals

We examined the fitted parameters of the model and compared these values across groups (Figure 3b). The discomfort data from all three groups is best fit by first scaling (α) the influence of melanopsin contrast by ∼60%. The scaled melanopsin and cone contrast is then combined with a sub-additive Minkowski exponent (β) of ∼1.75, intermediate between simple additivity (β=1), and a Euclidean distance metric (β=2). We find that these parameter values do not significantly differ between the three groups (Figure 3b, left). Therefore, we do not find that people with photophobic migraine differ from headache-free controls in the manner in which melanopsin and cone signals are combined at this initial stage.

The second pair of parameters convert log-transformed, ipRGC contrast into discomfort ratings. Here, substantial differences between the migraine and control groups were found. The MwoA group had a greater slope and a higher offset of discomfort rating, and the MwA group a higher offset, for a given amount of ipRGC contrast (Figure 3b, right). The migraine groups reported discomfort that was roughly twice as great overall and had a slope that was 50% steeper as compared to controls for the increase in discomfort with ipRGC contrast.

Based upon these results, we re-fit the model, forcing the stage 1 parameters to be the same across the three groups, but allowing the stage 2 parameters to vary. The output of stage 1 allows us to describe all the stimuli used in the experiment in terms of a single value of ipRGC contrast. Figure 3c re-plots the discomfort data for all participants and all stimuli from each group in terms of the stage 1 value of ipRGC contrast. The stage 2 model fits differ for each group and are used to generate the solid lines on the plots. Open circles mark the mean, across-participant discomfort ratings for each of the nine stimulus types. There is good agreement between the model fit and the across-participant mean discomfort. Forcing the stage 1 parameters to be the same across groups had minimal impact upon the fit of the model to the data (model R-squared, ±SEM: HAf controls: 0.95 ± 0.03; MwA: 0.96 ± 0.02; MwoA: 0.97 ± 0.01), supporting the claim that the stage 1 model parameters do not meaningfully differ between the groups.

Overall, these findings indicate that people with migraine with interictal photophobia do not differ from controls in the manner in which cone and melanopsin signals are scaled relative to each other and combined, but experience greater discomfort from this integrated signal.

### Migraine groups do not have enhanced pupil responses, indicating a selective enhancement of ipRGC discomfort signals

We considered the possibility that people with migraine have a general amplification of ipRGC signals at the level of the retina, of which visual discomfort is one aspect. If so, then we might expect an amplification of pupil responses to be seen in this population as well. To test this idea, we compared pupil constriction in the migraine groups to that observed in the headache-free control participants.

Figure 4a presents the mean, across-participant pupil responses observed in each of the three groups to the stimuli used in the experiment. The temporal profile of the pupil response to stimuli that target melanopsin, the cones, or their combination is in good agreement with prior reports^25^. There is also a clear increase in the amplitude of the pupil constriction produced by stimuli with increasing (100%, 200%, 400%) contrast.

**Figure 4.**
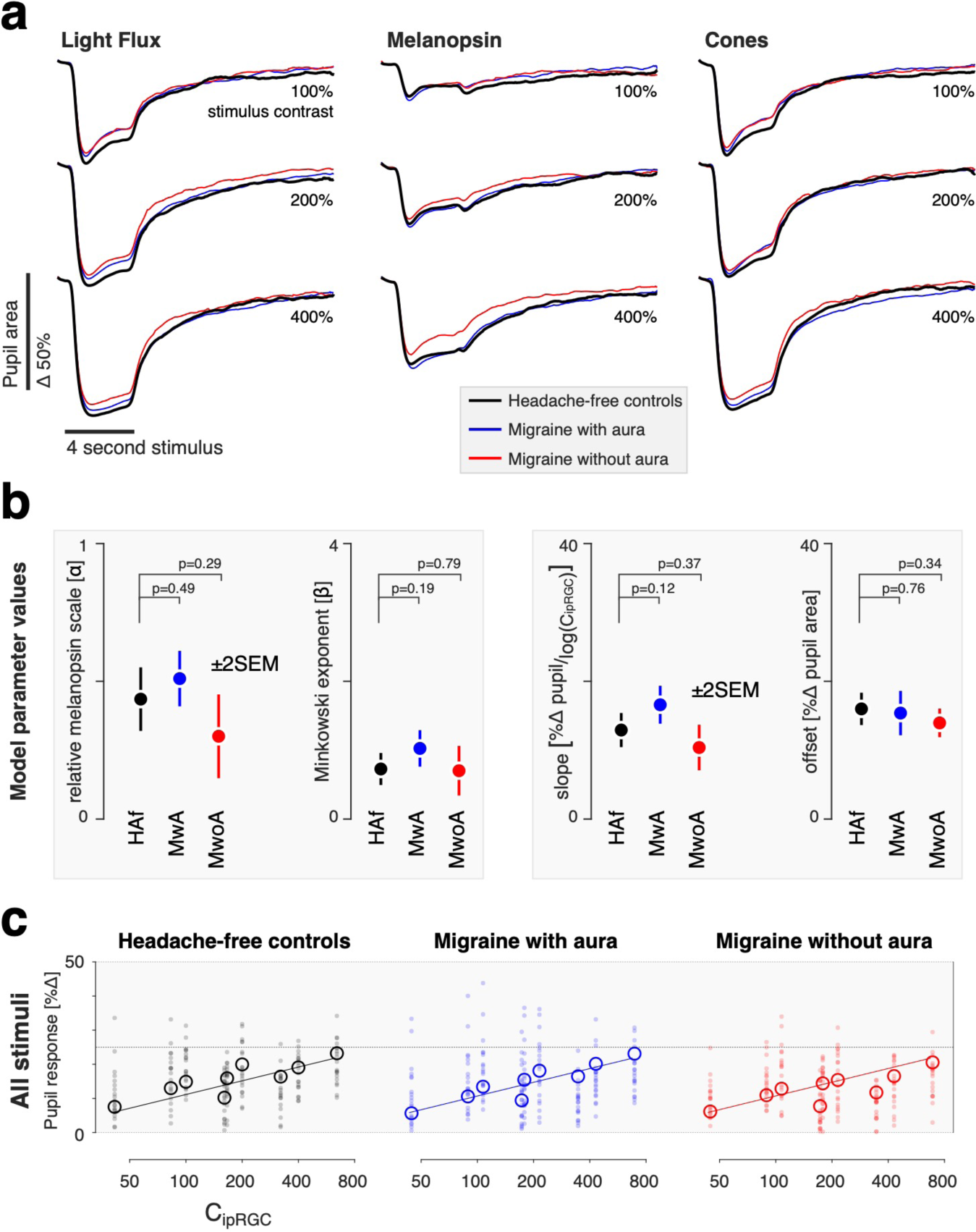
Pupil response by stimulus and group. **a**. The average pupil response across participants within each group (n = 20 participants per group) is shown for each stimulus type (columns) at each contrast level (rows). The responses from the three groups for each stimulus type are superimposed. **b**. We summarized the pupil responses by taking the mean of the percent change in amplitude of the pupil area across the recording period. These data were then fit with the two-stage model (see Figure 3). No significant differences between the groups in the parameter estimates were found (±2SEM obtained via bootstrapping), although both the relative melanopsin scaling and Minkowski exponent values are smaller for pupil responses than was observed for discomfort ratings. **c**. As no significant differences between groups was found for the parameters, we re-fit our model to the data forcing all parameters to be the same across groups. The plots report individual (filled circles) and mean (open circles) pupil response as a function of modeled ipRGC contrast.

The responses obtained from each studied group are close to overlapping in the plots for each combination of stimulus direction and contrast. We did not observe a greater amplitude of pupil response in the migraine groups as compared to the controls. Indeed, if anything, the pupil response in the migraine groups (particularly MwoA) is slightly attenuated compared to that of the headache free controls. We quantified the pupil response for each participant by measuring the mean percent change in pupil area following stimulus onset (Supplementary Figure 1). Similar to what was observed for visual discomfort ratings, the relationship between pupil response and stimulus contrast is well described as log-linear.

We next examined how cone and melanopsin signals are combined to produce the overall amplitude of pupil constriction, using the same two-stage model that we developed for the discomfort ratings (Figure 4b). The model fit the data from the three groups well (model R-squared, ±SEM: HAf controls: 0.95 ± 0.03; MwA: 0.98 ± 0.01; MwoA: 0.94 ± 0.02). We found no significant differences between the groups in the parameters of the model for pupil response. Therefore, we re-fit the model to the pupil data, forcing all parameters to be the same across the groups (Figure 4c). The agreement between the data and the model was quite good, despite requiring that all three groups be described using the same model parameters (model R-squared, ±SEM: HAf controls: 0.96 ± 0.01; MwA: 0.95 ± 0.001; MwoA: 0.91 ± 0.06). The stage 1 parameters in control of the pupil describe a scaling factor for melanopsin (α) of ∼40%, which is somewhat less than the influence that melanopsin has upon discomfort (∼60%). The Minkowski exponent for the combination of melanopsin and cone signals in the pupil response is ∼0.8, compared to its value of ∼1.75 for the discomfort ratings. The value of ∼0.8 indicates a combination rule for cone and melanopsin signals that is reasonably close to linear, consistent with prior observations of the additivity of cone and melanopsin signals in the pupil response^24,37^. The fact that the stage 1 parameters differ between the model fits to the two measures indicates that discomfort and pupil control are mediated by mechanisms that combine signals from melanopsin and the cones in different ways. A possible neural basis for these mechanisms would be distinct classes of ipRGCs.

Separately from the matter of how signals from melanopsin and the cones are combined across the two measures, the fact that the stage 2 parameters differ between controls and people with migraine for the production of discomfort but not for pupil constriction argues against the idea that a common, single amplification of retinal signals mediates increased interictal photophobia in migraine.

## Discussion

Our study indicates that the enhanced, interictal light sensitivity experienced by people with migraine is due to a selective amplification of a subset of ipRGC signals. Photophobia in migraine is not the result of an omnibus change in cone or melanopsin signals *per se*, but instead a change in the response to these photoreceptor inputs after they have been weighted and combined. Moreover, this amplification of retinal signaling is specific for discomfort signals, in that it is not observed for ipRGC outputs that control other reflexive responses to light, in particular pupil constriction.

### Distinct ipRGC classes

Studies in rodents^38–40^, primates^13,41–43^, and in the post-mortem human eye^44,45^ have demonstrated the existence of multiple classes and subclasses of ipRGCs, which differ in their photoreceptor inputs, signaling kinetics, and central projections. Control of circadian photoentrainment and the pupil response, for example, is attributable to distinct subsets of ipRGCs in rodents^38,46^.

We examined how melanopsin and cone inputs contribute separately and in combination to visual discomfort and to the pupil response within the context of a quantitative model. The first stage of our model estimates how melanopsin signals are weighted relative to cone signals, and the metric with which melanopsin and cone signals are combined. We did not find a difference at this stage between people with or without migraine. We did find, however, that the model parameters differ substantially when measured for the pupil response and for ratings of visual discomfort. A plausible explanation for these differences in photoreceptor combination is that different classes of ipRGCs contribute to visual discomfort and pupil responses in the human.

In the current study, we find that melanopsin and cone signals are combined approximately additively in control of the pupil, consistent with prior reports^24,37^. Melanopsin contrast was 40% as effective as cone contrast in modulating the pupil for these pulsed stimuli, as compared to a prior report of an overall 26% effectiveness of melanopsin relative to L+M cone modulations for driving pupil responses with sinusoidal modulations of contrast at low and high temporal frequencies^24^. We note that our index of pupil change here was across the entire time course of evoked response. While it is likely that the relative contribution of melanopsin to the amplitude of pupil constriction varies as a function of time following stimulus onset^19,25,37,47,48^, such a dissection of the components of the pupil response is beyond the scope of the current report.

In contrast to the pupil response, melanopsin and cone signals exhibit a nearly Euclidean combination metric in our measure of discomfort, and we find that the influence of melanopsin signals (relative to the cones) is ∼1.5x greater in producing visual discomfort as compared to pupil responses. A Euclidean combination metric is a feature of stimulus dimensions that produce a single, integrated percept^35^, suggesting that cone and melanopsin signals are combined into a unitary experience of discomfort.

We have previously found that observers describe targeted melanopsin stimulation as “uncomfortable brightness”^49^. It may be the case that the “brightness” and “discomfort” percepts, while each integrating cone and melanopsin signals, reflect the action of distinct retinal ganglion cell populations. Our present data, however, do not directly address such a dissociation.

Several studies have demonstrated that melanopsin contrast contributes to a sensation of brightness^50–53^. The melanopic component of brightness is presumably combined with the post-receptoral luminance channel that is derived from the sum of L and M cone excitations and carried by the classical (non-melanopsin containing) retinal ganglion cells. Yamakawa and colleagues measured the perceptual brightness of lights that varied in melanopic and luminance content^51^. A roughly additive effect of luminance and melanopsin content upon brightness is present in their data, although the form of the response departed from linear. The interpretation of these measurements is complicated, however, as the observers did not undergo pharmacologic dilation of the pupil, causing retinal irradiance to vary systematically with the stimulus.

Zele and colleagues also examined how cone and melanopsin signals combine in the perception of brightness^53^. Their work shows a log-linear relationship between isolated melanopsin and cone stimulus intensity and brightness. However, when presented in combination, they report two contribution components of cones to brightness, one of which is negative and may imply an adaptation process.

### Selective amplification

The ipRGCs are known to manifest linear changes in firing rates with logarithmic changes in retinal irradiance^54^. In our measurements, we find that ratings of visual discomfort, and the amplitude of evoked pupil response, vary linearly with log changes in stimulus contrast, consistent with an output system that receives these log-transformed signals from the ipRGCs.

While participant groups did not differ in the manner in which cone and melanopsin signals were combined, we find that people with episodic migraine with interictal photophobia have an amplification of the effect of this integrated signal upon ratings of visual discomfort. This amplification is similar in migraine with or without visual aura.

Importantly, we did not find evidence of a general amplification of ipRGC signals in migraine. The ipRGCs are the dominant, and perhaps exclusive, route for photoreceptor signals influencing the light-evoked pupil response via the pre-tectal nuclei^55,56^. If migraine is accompanied by a general amplification of ipRGC signals, then an enhanced light-evoked pupil response in this population might be predicted. Instead, we find that the amplitude of evoked pupil responses is not increased in people with migraine in response to stimulation of melanopsin, the cones, or their combination. Indeed, the trend in the data was towards smaller evoked pupil responses in migraine, especially in migraine without aura. Prior studies of pupil response in migraine have obtained varying results. Prior studies have not found migraine group differences in the amplitude of pupil constriction or steady-state pupil size^57–60^, although more subtle changes in pupil dynamics have been reported^58,61,62^. Our study differs from many prior reports in that we measured open-loop, consensual pupil responses by combining pharmacologic dilation with an artificial pupil, thus controlling retinal irradiance across the studied groups.

We also surveyed our participants regarding other behaviors that may be related to ipRGC function. A general alteration in ipRGC function in people with migraine might be predicted to be manifest in these measures as well. The ipRGCs have been implicated in circadian photoentrainment^15^, seasonal variation in mood and physiology^63–65^, and in the photic sneeze reflex. Our participants with migraine did not differ from headache-free controls in these behaviors, again suggesting that the amplification of ipRGC signals in migraine is specific to visual discomfort.

### The neural locus of amplification

While no specific ipRGC subtype has been identified as carrying the signal for visual discomfort, various lines of evidence implicate non-M1 ipRGCs^66–68^. The ipRGCs co-innervate neurons within the posterior thalamus of the rodent that also receive trigeminal afferents. These thalamic neurons then project onward to both somatosensory and visual cortices. Classes of ipRGCs also project to the lateral geniculate nucleus^13,69^ and are capable of modulating visual cortex responses^49^. Our findings of amplified discomfort to visual stimulation in people with migraine could reflect alteration of signals derived from the ipRGCs at any one of these sites.

A physiologic hallmark of migraine is alteration in the excitability of cortex, as manifest both in the phenomenon of cortical spreading depression of aura, and a tendency towards enhanced responses to sensory stimulation as compared to headache free controls^70^. Enhanced cortical responses to sensory stimulation has been observed in migraine with^70^ and possibly without^71^ aura, and for multiple sensory modalities. A natural locus, therefore, for the amplification of ipRGC signals for visual discomfort is at cortical sites. This could take place within primary visual or somatosensory cortex, or further downstream at the integration of these signals into a report of discomfort.

An ipRGC signal of visual discomfort might also be amplified at the level of the thalamus. Altered thalamic gating has been proposed as a mechanism for altered sensory perception in migraine, including photophobia^3,72^. Enhanced signaling within the trigeminal system may also contribute to amplification of ipRGC signals. In rodents, bright light activates the trigeminal ganglion and trigeminal nucleus caudalis^73–75^. Human studies suggest an interaction of the peripheral trigeminal system and light-mediated pathways, as noxious trigeminal stimulation lowers the visual discomfort threshold, and light stimulation lowers trigeminal pain thresholds^11,76^. Studies in the rodent implicate the ipRGCs in this interaction, as light aversion following corneal surface damage is attenuated in mice lacking ipRGCs^77^. Migraine may induce photophobia through the action of neuropeptides within this trigeminal-thalamic system^78^.

We might finally consider the possibility that ipRGC signals for visual discomfort are amplified at the level of the retina. This possibility strikes us as less plausible, given that our results would require a mechanism for selective enhancement of only the class of ipRGC that produces photophobia. Our results also argue against a change in the sensitivity of melanopsin or the cones in migraine under photopic conditions. There have been varying reports of alteration of cone electroretinogram responses in people with migaine^79,80^, although these studies are also difficult to interpret given possible differences in retinal irradiance between the studied groups^81^.

We interpret our results within a modeling approach that assumes that melanopsin and cone signals are integrated within the ipRGCs, and that post-retinal sites act upon the integrated, log-transformed signal. While this model was not a component of our pre-registered experimental protocol, we find that it provides an excellent account of the data. There is abundant evidence in support of the view that melanopsin and cone signals are integrated in the ipRGCs in this manner^13,21–23,82^. We cannot, however, exclude the possibility that cone and melanopsin signals are transmitted from the retina by separate channels, and that we are measuring the integration of these signals at some downstream site. Such a post-retinal integration is likely to be the case for the perception of the “brightness” of stimuli that combine melanopsin and cone contrast, as the post-retinal luminance channel originates in signals from the “classical” retinal ganglion cells, and must be integrated with signals from melanopsin-containing ipRGCs, perhaps at the level of the lateral geniculate nucleus.

More broadly, there is evidence that expression of melanopsin in eye tissues apart from the retina contributes to photophobia in rodent models^83^. Because we placed an artificial pupil between the stimulus and the pharmacologically dilated pupil of the observer, our stimuli illuminated only a small area of the cornea, and minimally the iris. There has also been interest in the contribution of the rods to photophobia in migraine^80^, and there is evidence that the rods provide inputs to ipRGCs^84^. We sought to minimize the influence of the rods upon our measurements by modulating our stimuli around a photopic background. While there is evidence that rod signals can modulate RGC firing at any light level^85^, the amplitude of these effects under photopic conditions is quite small relative to the cones. Further, our prior work indicates that rod signals do not make a measurable contribution to the pupil response at these background light levels^25^.

### Conclusion

Our study demonstrates that discomfort from light does not arise as the exclusive action of melanopsin, but instead reflects a signal that integrates cone and melanopsin inputs. The interictal photophobia of migraine is a selective amplification of this integrated signal, and one which does not extend to other domains of ipRGC function. We suspect that the amplification in migraine of ipRGC signals for discomfort occurs at a post-retinal site but cannot yet identify the locus. The modeling approach we adopted here provides a mechanism by which this localization might be pursued, by identifying central sites in which log changes in modeled ipRGC contrast are related to linear modulations of neural activity.

## Materials and Methods

We studied 20 participants in each of three groups: migraine with aura, migraine without aura, and headache-free controls (Table 1). Participants were between 25 and 40 years old and were recruited via digital social media. Headache classification was established using the Penn Online Evaluation of Migraine^86^. Participants with migraine were also required to endorse interictal photophobia^87^. Participants completed surveys that assessed behaviors putatively related to ipRGC function (Table 2).

Participants viewed stimuli that targeted specific photoreceptor classes using the technique of silent substitution^88^ (Figure 1c). Each stimulus type was presented at three, log-spaced contrast levels: 100%, 200%, and 400%. These stimuli were produced by a digital light synthesis engine (OneLight Spectra, Vancouver, BC, Canada) and tailored for the lens transmittance predicted for the age of each subject. The stimuli were presented through a custom-made eyepiece with a circular, uniform field of 27.5° diameter with the central 5° diameter of the field obscured to minimize macular stimulation. Spectroradiographic measurements were made before and after each session to ensure stimulus quality (Supplementary Table 1).

On each of many trials, the participant viewed a pulsed spectral modulation, at one of three contrast levels, designed to target melanopsin, the cones, or both (Figure 1c). The transition from the background to the stimulation spectrum (melanopsin, cones, or light flux), and the subsequent return to the background, were windowed with a 500 ms half-cosine. The total duration of the pulse was 4 seconds, after which the stimulus field returned to and remained at the background spectrum (Figure 1d). Twelve seconds after the pulse ended, participants were prompted by an auditory cue to verbally rate their visual discomfort on a 0-10 scale. Participants viewed the stimuli through their pharmacologically dilated right eye and a 6 mm diameter artificial pupil to control retinal irradiance. Infrared video recording of the left eye measured the consensual pupil response during each trial. Each participant viewed at least 12 trials for each crossing of photoreceptor target and contrast, and at least 6 of those trials were required to possess good quality pupillometry for the subject to be included in the study.

Pupil response was quantified for each trial as the mean percent change in pupil area during the period of 0 to 16 seconds from stimulus onset, relative to the 0.5 seconds before stimulus onset. We obtained the median pupil and discomfort response across trials within participant, and across participants within groups.

We examined the discomfort and pupil data within a two stage, non-linear model (Figure 3a). The response to a stimulus (either discomfort rating or pupil constriction) is given by:

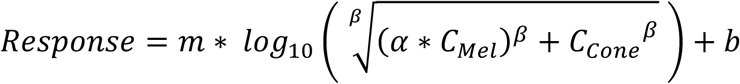

where C_Mel_ and C_Cone_ are the contrasts produced upon the melanopsin and cone photoreceptors by a stimulus, and *α, β, m*, and *b* are the four parameters of the model. Non-linear fitting was performed in MATLAB using *fmincon*, and the variability of parameter estimates within each group obtained by bootstrap resampling of the data across subjects.

This study was pre-registered (Supplementary Table 2). The analysis code is available (https://github.com/gkaguirrelab/melSquintAnalysis), as will be the data following publication.

Detailed methods are described in *SI Appendix, SI Text Online Methods*.

## Supplmental Material

### Supplementary Figures

**Supplementary Figure 1.**
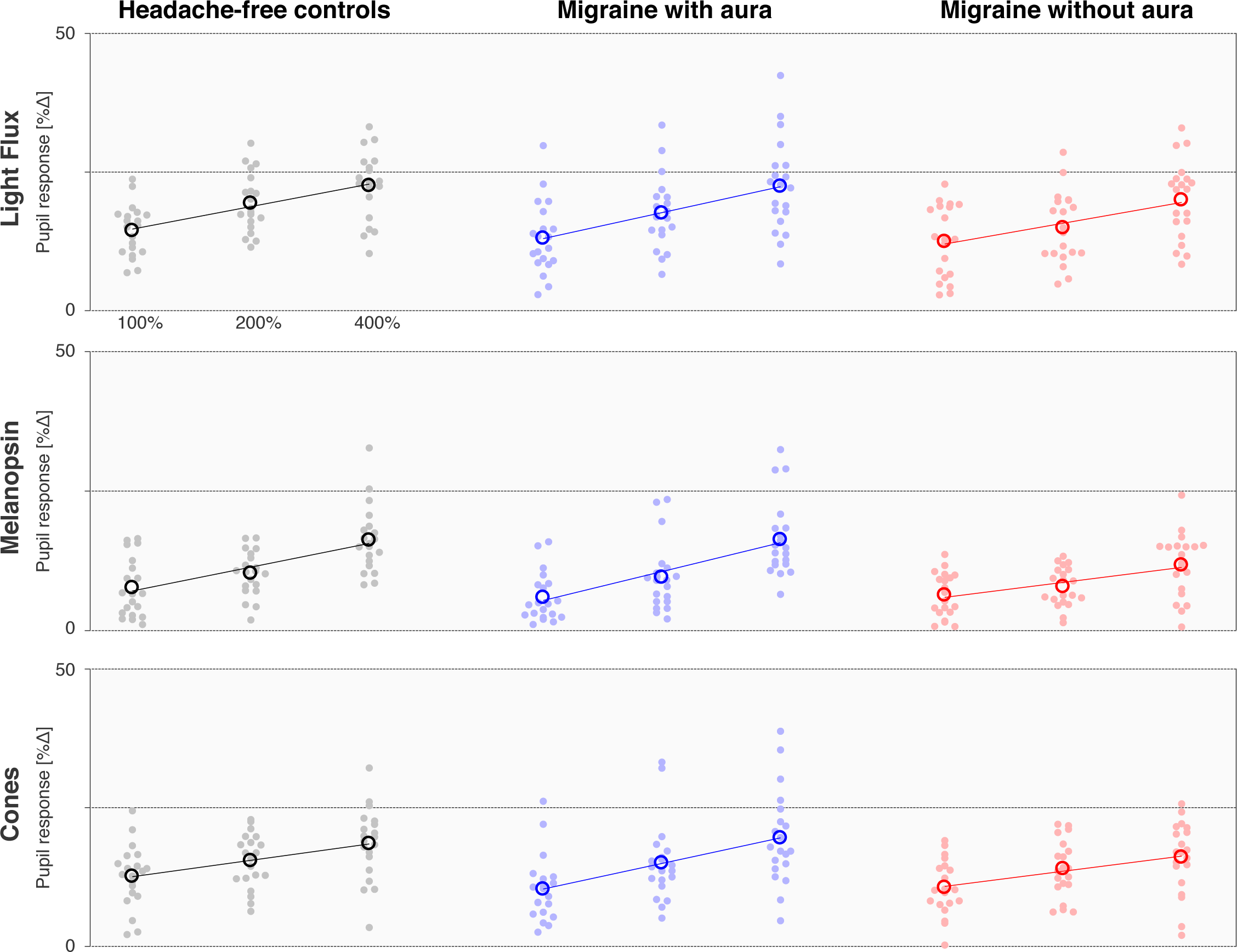
Pupil response amplitudes by stimulus and group. Each row presents the mean percent change in pupil area (across the duration of the trial) elicited by stimuli that targeted a particular combination of photoreceptors, and each column contains the data from each individual group (n = 20 participant per group). The stimuli were presented at three different contrast levels (100, 200, and 400%), and these values (log-spaced) define the x-axis of each subplot. The mean (across trial) pupil response for a given stimulus and contrast is shown for each participant (filled circle), as is the mean response across participants (open circle). The best fit line to the mean pupil response across participants as a function of log contrast is shown in each subplot.

### Supplementary Tables

**Supplementary Table 1.** Stimulus Validation Measurements. Before and after each experiment, we obtained 5 spectroradiographic measurements of each stimulus at its background and maximal (400% nominal) contrast level. From these measurements, we calculated the contrast on several receptor and post-receptoral mechanisms: melanopsin, the combined luminance channel (LMS), S-(L+M), and L-M. We also measured the luminance and chromaticity of the background spectrum for each stimulus. The upper portion of the table provides the mean of these validation measurements across sessions and subjects. The lower portion of the table presents the mean (across sessions and subjects) unsigned deviation of the validated stimuli from the nominal contrast levels.

| Stimulus | Contrast |  |  |  | Luminance | Chromaticity |
| --- | --- | --- | --- | --- | --- | --- |
|  | *Melanopsin* | *LMS* | *S* | *L-M* | [ cd/m2 ] | [ x, y ] |
| Mean stimulus values across subjects |  |  |  |  |  |  |
| *Mel* | 397.06% | 0.40% | -0.06% | -0.87% | 322 | 0.58, 0.38 |
| *Cones* | 0.83% | 397.72% | 1.72% | 1.02% | 145 | 0.57, 0.36 |
| *LF* | 398.74% | 399.41% | -2.09% | -0.08% | 209 | 0.57, 0.36 |
| Mean unsigned contrast error across subjects |  |  |  |  |  |  |
| *Mel* | 4.05% | 0.51% | 1.26% | 0.87% | - | - |
| *Cones* | 0.86% | 2.32% | 4.66% | 1.05% | - | - |
| *LF* | 1.42% | 1.27% | 3.33% | 0.41% | - | - |

**Supplementary Table 2.** Summary of Pre-registrations, addenda, and protocol deviations.

| Type | Link | Contents |
| --- | --- | --- |
| Initial pre-registration | https://osf.io/5ry67/ | - Defined initial plan for experiment, including subject recruitment, screening procedure, stimulus creation, experimental design, exclusion criteria, and primary analyses. |
| Addendum #1 | https://osf.io/qtjyd/ | - Modified subject exclusion criteria to exclude subjects with a history of ophthalmologic disease.<br>- Modified subject exclusion criteria to exclude candidate migraine participants who had not experienced a headache within the previous month. Seven migraine subjects with no headache in the prior month were enrolled in the study prior to the adoption of this criterion and are included in the resulting data set.<br>- Defined a specific target range for the luminance of the background stimulus spectrum.<br>- Added *'Morningness-Eveningness Questionnaire'*.<br>- Added a procedure to ask subjects in the week after a session if they have experienced a migraine since their participation in a session. |
| Addendum #2 | https://osf.io/kf253/ | - Defined a procedure to perform an examination of the pupil response data, masked to group assignment, to inform a revision of subject recruitment targets. |
| Addendum #3 | https://osf.io/j3x24/ | - Revised recruitment goal to 20 subjects within each group from the original target of 40. |
| Deviations |  | - We had proposed to report median values across subjects. Using the median value result in unstable bootstrap model fits due to the discontinuous nature of the discomfort ratings data. We therefore switched to the mean for all across-subject measures. |

### Supplemental Methods

We used silent substitution to create stimuli designed to selectively target melanopsin, the cones, or both (Figure 1c). We presented 4 second pulses of these stimuli to participants and asked them to verbally report the degree of discomfort they experienced while simultaneous pupillometry was recorded (Figure 1d,e).

The study was approved by the Institutional Review Board of the University of Pennsylvania. All participants provided informed written consent, and all experiments adhered to the tenets of the Declaration of Helsinki.

#### Participants

A total of 60 participants between the ages of 25 and 40 were recruited from the greater Philadelphia area and University of Pennsylvania campus, in most cases using advertising on digital social media services. All candidate participants underwent screening using the Penn Online Evaluation of Migraine^1^, which implements automated headache symptom classification using the International Classification of Headache Disorders (ICHD)-3-beta criteria. The POEM also incorporates a set of previously published questions regarding photophobia during and between headache. These responses were scored with a point for each yes response to questions 1 through 7 (referred to here as the Choi score)^2^. Potential participants also completed the Visual Discomfort Score (VDS) survey^3^. The VDS score was derived as the sum of scores from 23 questions regarding frequency of particular visual discomfort symptoms, each scored on a 0-3 scale from “never” to “almost always”. To be eligible for the study, potential participants were required to meet all inclusion criteria for one of three groups:

1. Migraine with visual aura (MwA): a) classification of migraine with visual aura by the POEM, b) Choi score of 6 or 7, c) a response of “yes” to the Choi query regarding the presence of photophobia during headache free periods, d) one or more headaches within the prior month.
2. Migraine without aura (MwoA): a) classification of migraine without aura by the POEM, b) Choi score of 6 or 7, c) a response of “yes” to the Choi query regarding the presence of photophobia during headache free periods, d) one or more headaches within the prior month.
3. Control: a) classification of mild non-migrainous headache or headache-free by the POEM, b) a response of “No” or “I don’t know” to a question regarding a family history of migraine, c) a response of “No” to a question regarding a history of childhood motion sickness, d) VDS score 7 or lower.

Exclusion criteria were a history of glaucoma, generalized epilepsy, a history of adverse reaction to dilating eye drops, a concussion in the last six months, ongoing symptoms from head trauma/concussion, best-corrected distance acuity below 20/40 assessed via Snellen eye chart, or abnormal color vision as assessed via Ishihara plates. Participants were not excluded based on medication use, including migraine preventive medications, and were allowed to continue to take their current medications during data collection.

An inability to collect usable pupillometry from a participant was an additional exclusion criterion. Candidate participants were studied in the lab during a pupillometry screening session that mimicked a subset of trials from the main experiment. To pass this screening session, participants were required to provide acceptable pupillometry data on at least 9 of 12 trials. We screened 83 participants, and 2 were excluded on the basis of this screening criterion. Drawing from the 81 participants who met the pupillometry screening criterion, we ultimately collected archival data on 60 participants, with 20 participants from each group (MwA, MwoA, and controls). Of the remaining 21 participants, 15 either did not respond to subsequent attempts to enroll in the study or were screened after data collection had completed. An additional 6 participants participated in at least one session but failed to return for subsequent sessions.

#### Stimuli

We designed stimuli that target specific photoreceptor classes through the technique of silent substitution^4^. We targeted three main photoreceptor mechanisms: melanopsin, the cones, or both (Figure 1c). We use here the term “light flux” to describe the latter combined stimulus, although we note that our stimulus modulation is not simply a multiplicative scaling of the spectrum, which is what is usually implied by the term.

These stimuli were generated in the same manner as described in prior reports^5,6^. Briefly, we used a digital light synthesis engine (OneLight Spectra, Vancouver, BC, Canada) that produces stimulus spectra as mixtures of 56 independent primaries (∼16 nm full width at half maximum) under digital control and can modulate between these spectra at 256 Hz. We tailored the photoreceptor spectral sensitivities for each individual observer, taking into account the participant’s age, pupil size, and our field size of 27.5 degrees (Figure 1f). Cone fundamentals were based on the International Commission on Illumination (CIE) physiological cone fundamentals^7^. The CIE standard only specifies fundamentals up to field sizes of 10 degrees, so we obtain estimates for our 27.5 degree field by extrapolating the formula using the open-source Psychophysics Toolbox^8–10^. We created separate background and stimulation spectra that provided 1) a nominal 400% unipolar Weber contrast on melanopsin while silencing the cones for our melanopsin-directed background/stimulus pair (melanopsin direction), 2) 400% contrast on each L-, M-, and S-cone classes while silencing melanopsin for the cone-directed modulation/stimulus pair (cones direction), and 3) 400% contrast each on melanopsin and each L-, M-, and S-cone classes for the light flux modulation/stimulus pair (light flux direction) (Figure 1c). The background spectra for each stimulus type differed in background luminance, but had similar chromaticity (Supplementary Table 1) as calculated using the XYZ functions associated with the CIE 2006 10-degree cone fundamentals (http://www.crvl.org)^7^. We also produced contrast levels of 100 and 200% for each stimulus direction by scaling the relevant stimulus spectrum. We did not explicitly silence rods or penumbral cones^6^, although we believe that the luminance and temporal properties of our stimuli minimize the contribution of these photoreceptors.

Stimuli were presented through a custom-made eyepiece with a circular, uniform field of 27.5° diameter. The central 5° diameter of the field was obscured to block the effects of the foveal macular pigment which can cause variation in photoreceptor spectral sensitivity (Figure 1f).

We obtained 5 spectroradiometric measurements of the 400% stimuli and their backgrounds before and after each testing session. For each stimulus type, we determined contrast on the following post-receptoral mechanisms: LMS, L-M, S, and melanopsin. Supplementary Table 1 presents the average validation results across all sessions. The validation results for each session are included with the experimental data for download. We adjusted our apparatus over the duration of data collection to maintain the luminance of the background spectrum for the light flux stimulus between 160 and 254 cd/m^2^.

We discarded data from a session if the post-experiment stimulus validation measurements did not meet our pre-registered criteria. Data were discarded if the median of the 5 post-experiment measurements was: 1) greater than 20% absolute contrast on any of the nominally silenced post-receptoral mechanisms; or 2) less than 350% contrast upon a targeted post-receptoral mechanism. In the event that data from a session were discarded, the session was repeated at a later date. Prior to starting a data collection session, pre-experiment stimulus validation measurements were required to meet these same criteria.

#### Experiment Structure

Each participant was studied during multiple data collection sessions, usually held on different days. In an attempt to minimize variation in circadian cycle across sessions, subsequent sessions were initiated within three hours of the time of day when the same participant started their first session.

Participants were exposed to similar “light history” prior to data collection. At the start of a session, participants entered the testing room and underwent pharmacologic dilation of their right eye with 0.5% proparacaine for anesthesia and then 1% tropicamide ophthalmic solution. Participants remained in the testing room for the next 20 minutes, receiving instructions and adjusting the position of the apparatus for comfort. Room lights were set so that the walls of the testing room had a measured luminance of ∼150 cd/m2, equated to the background luminance of our light flux stimulus. After confirming the presence of pupil dilation, the room lights were turned off and a curtain closed behind the participant to block light from the screen of the computer that controlled the apparatus. Apart from the light from the eyepiece, the participant remained in darkness for the remainder of the experiment. Participants viewed the stimuli through their pharmacologically dilated right eye and a 6 mm diameter artificial pupil to control retinal irradiance.

On each of many trials, the participant viewed a pulsed spectral modulation designed to target melanopsin, the cones, or both (Figure 1c). The transition from the background to the stimulation spectrum (melanopsin, cones, or light flux) and the subsequent return to the background were windowed with a 500 ms half-cosine. This step minimized the entopic percept of a Purkinje tree in the melanopsin-directed stimulus^6^. The total duration of the pulse was 4 seconds, after which the stimulus field returned to and remained at the background spectrum (Figure 1d). Twelve seconds after the pulse ended, participants were prompted by an auditory cue to verbally report their discomfort rating (described below). This verbal rating was recorded by a microphone during the 4 second response window, the end of which was marked by another auditory cue. There was a variable inter-trial-interval of 1.5 – 2.5 seconds (uniformly distributed) that reduced the predictability of the onset of the next trial.

Ten consecutive trials that targeted the same photoreceptor direction but varied in contrast were grouped together into an acquisition. The ordering of the contrast levels (100, 200, 400%) followed a counterbalanced sequence to avoid trial order effects^11^; the first trial was discarded so that all retained trials had controlled first-order stimulus history. A total of 6 acquisitions, 2 of each stimulus type, comprised a single session. Acquisitions were ordered such that consecutive acquisitions were not of the same stimulus class. Data collection for a participant was deemed sufficient when a subject had completed two sessions, and these sessions contained in total at least six acceptable trials—as judged by pupillometry—for each stimulus type. Only acceptable trials were included in pupillometry analysis, but all trials were included in analysis of discomfort ratings. We attempted to gather 4 sessions of data for each individual participant but retained all subjects with data collection that was deemed sufficient. Participants did not complete all 4 sessions for a variety of reasons, including failure of post-experiment stimulus validation, poor pupillometry requiring us to discard that session, or declining to return for subsequent sessions. Across all 60 participants, 45 completed 4 sessions (15 controls, 15 MwA, 15 MwoA), 11 completed 3 sessions (4 controls, 3 MwA, 4 MwoA), and 4 completed 2 sessions (1 control, 2 MwA, 1 MwoA).

#### Discomfort Ratings

At the end of each trial, participants were asked to rate the discomfort produced by the stimulus on a 0 – 10 scale. The experimenter read this prompt to the participant at the start of each session:

> “Following each trial, please rate the degree of discomfort that you experienced from the light pulse on a scale of zero to ten. A score of zero means that the pulse was not at all uncomfortable. A score of five means that the light pulse was moderately uncomfortable. A score of ten means that the light pulse was extremely uncomfortable.”

Following completion of the experiment, raters masked to group assignment manually transcribed these verbal discomfort ratings. Trials on which no rating was given, or on which the spoken rating was un-interpretable, were discarded.

#### Pupillometry

We recorded the consensual pupil response from the left eye of the participant (contralateral to the eye receiving the stimulus) using an infrared camera (Pupil Labs GmbH) mounted on a post ∼25 mm from the eye. A video clip was recorded for each trial, starting 1.5-s prior to pulse onset and ending 12-s after pulse offset (Figure 1d,e). These videos were processed using custom software (https://github.com/gkaguirrelab/transparentTrack)^12^ to fit an ellipse to the identified pupil boundary in each video frame, allowing us to extract pupil area over time.

This raw pupillometry data underwent several stages of pre-processing to remove and interpolate over frames in which the pupil had been poorly identified. The first stage involves blink censoring, which was performed by identifying frames in which the glint from the active infrared light source of the camera was absent. Several frames before and after each blink were also censored to remove blink-related artifacts, with these values adjusted on a per-participant basis. Next, frames in which the pupil was identified but poorly fit by the routine were censored. This step largely functioned to remove frames in which much of the pupil was obscured by the eyelid. Lastly, frames in which the identified pupil was implausibly large or small were removed. Linear interpolation was performed over censored frames. If more than 25% of frames in a given trial were censored that trial was discarded from analysis. Pupil responses were expressed as the percentage change in area relative to the 0.5-s prior to the stimulus onset.

Six participants had frequent, brief blinks that produced many missed frames of pupil tracking despite having video recording of the eye that was otherwise of good quality. The data from these participants were retained despite having more than 25% missing frames in a trial.

All manual adjustment of pupillometry, and indeed the development of the processing steps and criteria, was performed by investigators masked to the group membership of the participants.

#### Analysis

Data analysis was performed using custom MATLAB code (Mathworks). We used a one-way ANOVA to determine the effect of group upon the clinical and demographic measures. Significant effects were examined in post-hoc testing using the Tukey procedure.

We took the median of discomfort rating across trials within a participant, and the mean across participants within a group. Pupil response was quantified for each trial as the mean percent change in pupil area during the period of 0 to 16 seconds from stimulus onset, relative to the 0.5 seconds before stimulus onset. The mean response across trials within participant, and across participants within groups, was obtained.

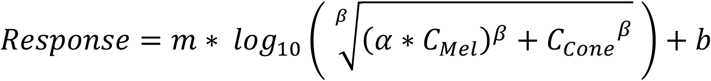

where C_Mel_ and C_Cone_ are the contrasts produced upon the melanopsin and cone photoreceptors by a stimulus, and *α, β, m*, and *b* are the four parameters of the model.

The first two model parameters describe an initial, non-linear stage that combines the photoreceptor contrast inputs and provides an “ipRGC contrast” output. The melanopsin contrast (*α*) is weighted by the first parameter, and then the weighted melanopsin contrast and the cone contrast are then combined using the Minkowski distance metric, with the second parameter (*β*) being the exponent for the metric. The modeled ipRGC contrasts of the stimuli are then base-10 log transformed and passed through a two-parameter affine transformation (parameters, slope and intercept of the transformation). In reporting the parameters, we convert the intercept parameter to an “offset” value, which is the predicted response amplitude at an ipRGC contrast of 200%.

The model was fit to the mean (across participant) data for all stimuli, with separate models fits performed for each group, and for each data type (discomfort rating and pupil response). Model fitting was performed using the *fmincon* function in MATLAB to minimize the L2 norm between the modeled values and the data. This fitting procedure was repeated over 1000 bootstrap resamples (with replacement) of the participants to assess the variability of the model output. We observed that the *β* parameter deviated slightly from a normal distribution across bootstraps. Therefore, we obtained the median value for all parameters across bootstraps, and variability across bootstraps was expressed by dividing the inter-quartile range of the values across bootstraps by 1.35, yielding a measure commensurate with the standard deviation of the distribution and thus an estimate of the standard error of the mean of the central tendency of the parameters.

#### Pre-Registration and availability of data and analysis code

This study was the subject of an initial pre-registration document (https://osf.io/5ry67/) and subsequent addenda (project summary page: https://osf.io/qjxdf/). Supplementary Table 2 summarizes the pre-registration documents and our deviations from these protocols. The analysis code is available (https://github.com/gkaguirrelab/melSquintAnalysis), as will be the data following publication.

